# Comparison of seven single cell Whole Genome Amplification commercial kits using targeted sequencing

**DOI:** 10.1101/186940

**Authors:** Tamir Biezuner, Ofir Raz, Shiran Amir, Lilach Milo, Rivka Adar, Yael Fried, Elena Ainbinder, Ehud Shapiro

**Affiliations:** Department of Computer Science and Applied Mathematics, Weizmann Institute of Science, Rehovot 761001, Israel; Stem Cell Core and Advanced Cell Technologies, Life Sciences Core Facilities, Weizmann Institute of Science, Rehovot 761001, Israel

## Abstract

Advances in biochemical technologies have led to a boost in the field of single cell genomics. Observation of the genome at a single cell resolution is currently achieved by pre-amplification using whole genome amplification (WGA) techniques that differ by their biochemical aspects and as a result by biased amplification of the original molecule. Several comparisons between commercially available single cell dedicated WGA kits (scWGA) were performed, however, these comparisons are costly, were only performed on selected scWGA kit and more notably, are limited by the number of analyzed cells, making them limited for reproducibility analysis. We benchmarked an economical assay to compare all commercially available scWGA kits that is based on targeted sequencing of thousands of genomic regions, including highly mutable genomic regions (microsatellites), from a large cohort of human single cells (125 cells in total). Using this approach, we could analyze the genome coverage, the reproducibility of genome coverage and the error rate of each kit. Our experimental design provides an affordable and reliable comparative assay that simulates a real single cell experiment. Results demonstrate the need for a dedicated kit selection depending on the desired single cell assay.

## Introduction

The increase in throughput and precision of next generation sequencing (NGS) in recent years has adramatic effect on biological research. Cell to cell variability within the same organism became a highly investigated research study, underlying the need for new and improved molecular biology tools for the purpose of accurate analysis of various single cell (SC) properties (*e.g.* gene expression, genomics, epigenomics) in a scalable manner^1^. SC genome variability is a fascinating example for the need for accurate measurements as sequence changes (*e.g.*somatic mutations, cancer driven mutations) variations occur during development in health (*e.g.* random somatic mutations, genomic recombination during B and T cell maturation) and disease (*e.g.* driver mutations and copy number alterations in cancer). Moreover, SC genomics has also extended to capture more than one attribute of a cell, emphasizing the need for an integrated multi-omics SC analysis^2^.

Since single molecule sequencing is still in its early stages, a variety of Whole Genome Amplification (WGA) protocols, which amplify the entire genome are the current state-of-the-art in SC genome analysis. A genome contains a single copy of each nucleotide and hence, any biased part of the analysis pipeline, both biochemical and both computational, that leads to modification or loss of information will have a dramatic effect on the conclusion of an experiment^3^. Examples for biased amplification are: *in vitro* mutation insertion/deletion, loss of genomic regions (allelic drop out-ADO) and non-uniform amplification (that leads to incorrect copy number variation analysis). The reproducibility of the protocol is sometimes even more important than the above, for example when SCs are to be compared between their sequences, high genome coverage can be less effective than the reproductive amplification of the same sequences in every cell, such that their data can be intersected during analysis^4^.

WGA protocols differ by various parameters, namely by the polymerase type, and the molecular biology principle standing behind the amplification (as reviewed here:^3,5^). WGA protocols originally emerged to enable the analysis of low starting DNA material, and in recent years single cell dedicated WGA kits (scWGA) have emerged and commercialized, to enable accurate amplification at a SC resolution, starting from ~6pg of DNA. For this comparison all 7 commercially available SC dedicated WGA kits (all kits known to the authors until the date of the experiment design) were selected. Although scWGA kit comparisons were published^6^^-^^13^, none has yet to compare all of the available kits in a single comparison, and at most, selected kits that represent the same category were selected for comparison. Additionally, some are based on non-NGS based analysis^10^ and those that do, either sequence non-eukaryotic cells^6^ or are limited by the number of cells per kit (<9 cells, and in some cases only 2-3 cells)^7-9,11,13^, affecting the reproducibility ofthe results. The high costs involved in SC genomics, which include the cost of a reaction of scWGA reaction and costs of downstream analyses (*e.g.* whole exome sequencing (WES) or whole genome sequencing (WGS) of many cells) reason the lack of large-scale comparison experiments.

The goal of this study is to compare all of the available scWGA kits by using a previously established targeted enrichment approach - using a highly mutltiplexed microfluidics chip Access Array, Fluidigm^14^. Specifically, single cells, ranging between 12 to 23 cells per scWGA kit (125 single cells in total) were analyzed by a PCR panel of 3401 amplicons, mainly comprised Microsatellite (MS) containing amplicons 3233 (95% of the panel). Taking advantage of the large cohort of cells as a means for increased reliability of the results (similar to a real experiment), sequencing analysis aimed to compare amplification bias properties per each scWGA kit. The following aspects were analyzed: genome coverage, reproducibility of amplification between single cells (intersecting successfully amplified loci) and, due to the instable nature of MS synthesis *in vitro*, the error-rate of each scWGA kit.

## Results

### Generation of a large cohort of single cells data for scWGA kits comparison

In order to create a comprehensive analysis of scWGA kits, we have approached the companies/vendors of all currently available scWGA kits (Table 1). All companies replied and thankfully delivered reagents for the comparison experiment. Following a preliminary experiment which provided evidence that deposition of SCs using the CellCelector (ALS) cell picker to a <5μl deposition buffer is technically impractical (relevant to Ampli1, Genomphi, and TruePrime, data not shown), appropriate kit providers were approached to supply a working protocol for 5μl deposition volume (see Methods).

**Table 1.**
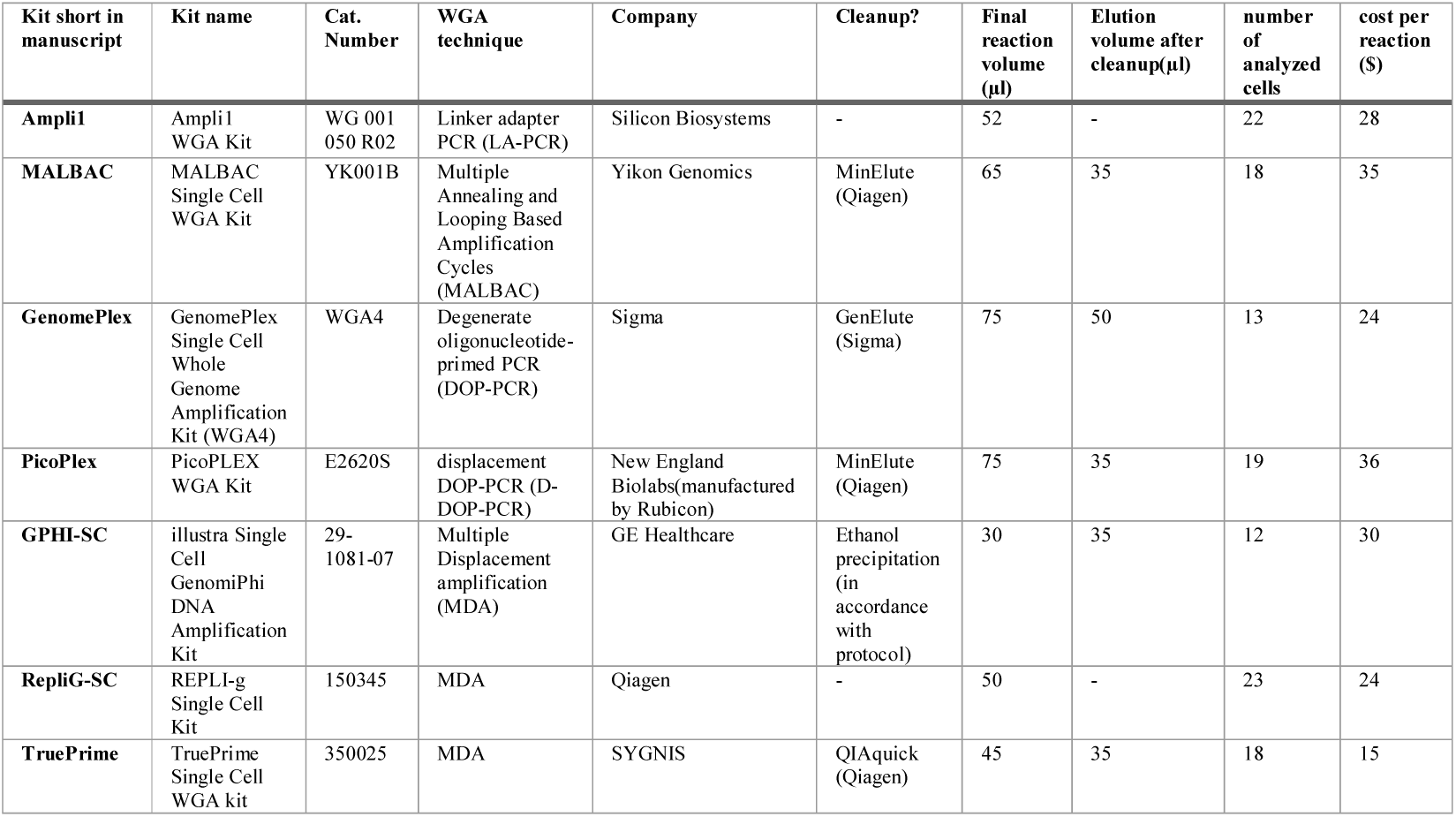
Summary of participating scWGA kits. We first created a clone from a single human ES cell (H1) in order to create a uniform population of normal cells (without known chromosomal aberrations). Before cell picking we have prepared a dedicated 96 well PCR plate for each scWGA kit containing the appropriate deposition reagent in each reaction well (PBS or specific lysis buffer). Using an automated cell picking device (CellCelector, ALS) we picked and deposited SCs to each reaction well of the scWGA kit dedicated 96 well plate. scWGA reactions were immediately proceeded in accordance with manufactures’ protocols. Following WGA process, DNA samples were randomized and processed by our lab developed streamlined cell lineage analysis platform using Access Array (AA, Fluidigm) chips to retrieve targeted enrichment of 3401 amplicons (Supplementary Table 1). 95% of the amplicons in the panel comprise of 4282 MS loci, as some amplicons contain more than a single MS locus (Supplementary Table 2). 74% of MS containing amplicons target the X chromosome, to enable an uninterrupted analysis of genome coverage in asingle allele occurrence in male normal cells. Following NGS data mapping to the corresponding panel amplicons, a summary table was composed, counting the number of mapped reads for each amplicon in each DNA sample (Supplementary Table 3). To validate the robustness and accuracy of the platform, we first analyzed: (a) the reproducibility of the results for negative and positive controls (di-distilled water (DDW) and Hela genomic DNA, respectively) deposited in each of the five participating AA chip (5 samples for each control). For that we measured the success rate of the read mapping (# mappedreads/Total reads per sample) and by counting the # of mapped loci from the entire panel (Supplementary Figure 1a and b, respectively). (b) the replication of the same analyses as in (a) for 40 duplicates: 39 single cell duplicates and one bulk DNA from the H1 cell line duplication), distributed randomly across the participating AA chips (Supplementary Figure 1c and d, respectively). One of the features of the platform is the use of the Echo550 (Labcyte) to gain an equal representation of the samples in a highly multiplexed NGS run following a low coverage Miseq run. We have normalized the sample volumes for a NextSeq run, such that all samples will have an equal read count (besides failed samples that were given a chance by pooling them at a fixed volume). The Interquartile Range (IQR) of the percentage of total reads per sample as part of the sum of all reads was reduced from 0.6% in the Miseq run to 0.1% in the NextSeq run. This normalization enabled a comparable analysis of all scWGA kits as it will perform on a real experiment, as a low success rate means that more money is being spent on non-informative reads. For further analyses, only single cells were used (unless specifically mentioned). A single replicate sample was randomly selected from each cell sample duplicates (mentioned in Supplementary Table 3).

| Kit short in manuscript | Kit name | Cat. Number | WGA technique | Company | Cleanup? | Final reaction volume (µl) | Elution volume after cleanup(µl) | number of analyzed cells | cost per reaction (\$) |
| --- | --- | --- | --- | --- | --- | --- | --- | --- | --- |
| **Ampli1** | Ampli1 WGA Kit | WG 001 050 R02 | Linker adapter PCR (LA-PCR) | Silicon Biosystems | - | 52 | - | 22 | 28 |
| **MALBAC** | MALBAC Single Cell WGA Kit | YK001B | Multiple Annealing and Looping Based Amplification Cycles (MALBAC) | Yikon Genomics | MinElute (Qiagen) | 65 | 35 | 18 | 35 |
| **GenomePlex** | GenomePlex Single Cell Whole Genome Amplification Kit (WGA4) | WGA4 | Degenerate oligonucleotide-primed PCR (DOP-PCR) | Sigma | GenElute (Sigma) | 75 | 50 | 13 | 24 |
| **PicoPlex** | PicoPLEX WGA Kit | E2620S | displacement DOP-PCR (D-DOP-PCR) | New England Biolabs(manufactured by Rubicon) | MinElute (Qiagen) | 75 | 35 | 19 | 36 |
| **GPHI-SC** | illustra Single Cell GenomiPhi DNA Amplification Kit | 29-1081-07 | Multiple Displacement amplification (MDA) | GE Healthcare | Ethanol precipitation (in accordance with protocol) | 30 | 35 | 12 | 30 |
| **RepliG-SC** | REPLI-g Single Cell Kit | 150345 | MDA | Qiagen | - | 50 | - | 23 | 24 |
| **TruePrime** | TruePrime Single Cell WGA kit | 350025 | MDA | SYGNIS | QIAquick (Qiagen) | 45 | 35 | 18 | 15 |

### scWGA genome coverage analysis

One of the key measurements in SC genome research is the percentage of the genome that yields from the amplification. Using our amplicon panel, we sought to validate the genome coverage by counting the number of successful amplicons (>0 reads). Improper amplification may lead to misinterpretation of the data for genome coverage analysis, for example, if a single allele will not be amplified, it might be undetected as the other allele may “compensates” for the loss. To tackle this we counted only amplicons on the X chromosome (H1 is a normal diploid male cell line) and used H1 cell line bulk preparation as a control (2 duplicates of the same bulk extraction). An inherit bias in Ampli1 kit is that it starts with a biased digestion of the genome (MseI - sequence recognition site“TTAA”). Hence, a fair comparison between kits would exclude all amplicons that include MseI recognition regions (see biased Ampli1 results in Supplementary Figure 2). Plotting all single cells per kit in a single graph (Figure 1) shows that Ampli1, and later RepliG-SC are the best at the genome coverage aspect, yielding medians of 1095.5 and 918 amplicons per single cell, respectively. Best amplifying cells in the experiment are coming from these two kits. GPHI-SC, PicoPlex, and MALBAC are next in this category with medians of 807.5, 750 and 696.5 amplified loci per single cell, respectively. Notably, PicoPlex is the most reliable kit, with the tightest inter quartile region (IQR) of all kits, and no failed cell. Specific experimental calibrations may assist in reducing the failed cells (improvements of picking, modifying the protocols etc.). GenomePlex and TruePrime generated significantly less unmapped reads (Supplementary Table 3) and a low number of mapped amplicons. Their positive control results are similar to the SC data, leading to the conclusion that the poor results are not due to cell picking procedure but a genuine failure of kit performance.

**Figure 1.**
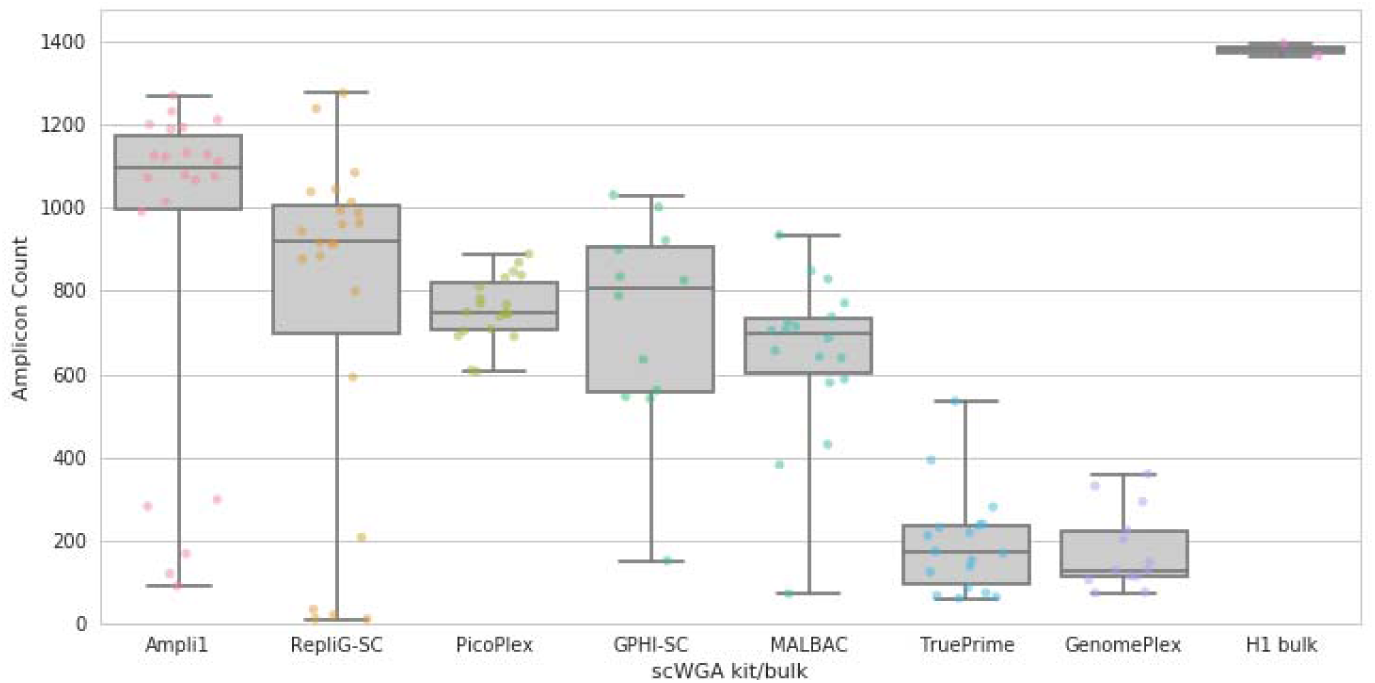
Amplicon coverage per single cell kit of only “TTAA” free amplicons. Mapped amplicons were counted per each single cell. Data represents only MseI restriction site free amplicons (“TTAA”), to follow the internal bias of the Ampli1 against these amplicons (see Supplementary Figure 2). Each dot represents a single cell, except for the right column, where each dot represents a cell bulk duplicate, originated from the same cell line (H1). Each column is the collection of all single cells per scWGA kit (except for the H1 bulk column).

### scWGA reproducibility analysis

In some cases, the reproducibility of the scWGA kit over several SC samples is more important than other important parameters (such as a genome coverage). Having a large number of cells allows for a more reliable understanding of a real experimental reproducibility, taking into account all cell population, including unsuccessful SC WGA product. In order to analyze reproducibility, we have generated a dataset of all possible groups of two cells per kit and counted the number of intersecting loci forall calculated groups, with the restriction of MseI regions, as explained above (Figure 2a). Results show that Ampli1 is the most reproducible kit, amplifying more loci than its follower, RepliG-SC. Notably, we can see clusters of cell groups that partially or completely failed. To get a better simulation of a real experiment, where successfully amplified cells are selected for analysis, we have selected the cells from the upper median of the most successful cells (Figure 2b) as reflected by their number of amplifying loci, as presented in Figure 1 (all cells above median amplicon count), and repeated the analysis. As expected, results show a tighter range of intersecting amplicons, again, showing the better reproducibility of Ampli1 amplification. A repeat of the same type of analysis for groups sizes of k = 3 and 4 cells shows similar results (Figure 2c and d), with an expected drop of intersecting amplicons as cell group size increases. Interestingly, analysis shows a mild decrease in the number of intersecting loci as increasing the group size for all kits. This provides a strong evidence that WGA protocols are systematically biased for the loci they amplify from the genome. PicoPlex, although not the best in the aspect of number of amplifying loci when compared to other kits, demonstrates high reproducibility for all of its cells (Figure 2c) supporting the biased amplification assumption.

**Figure 2.**
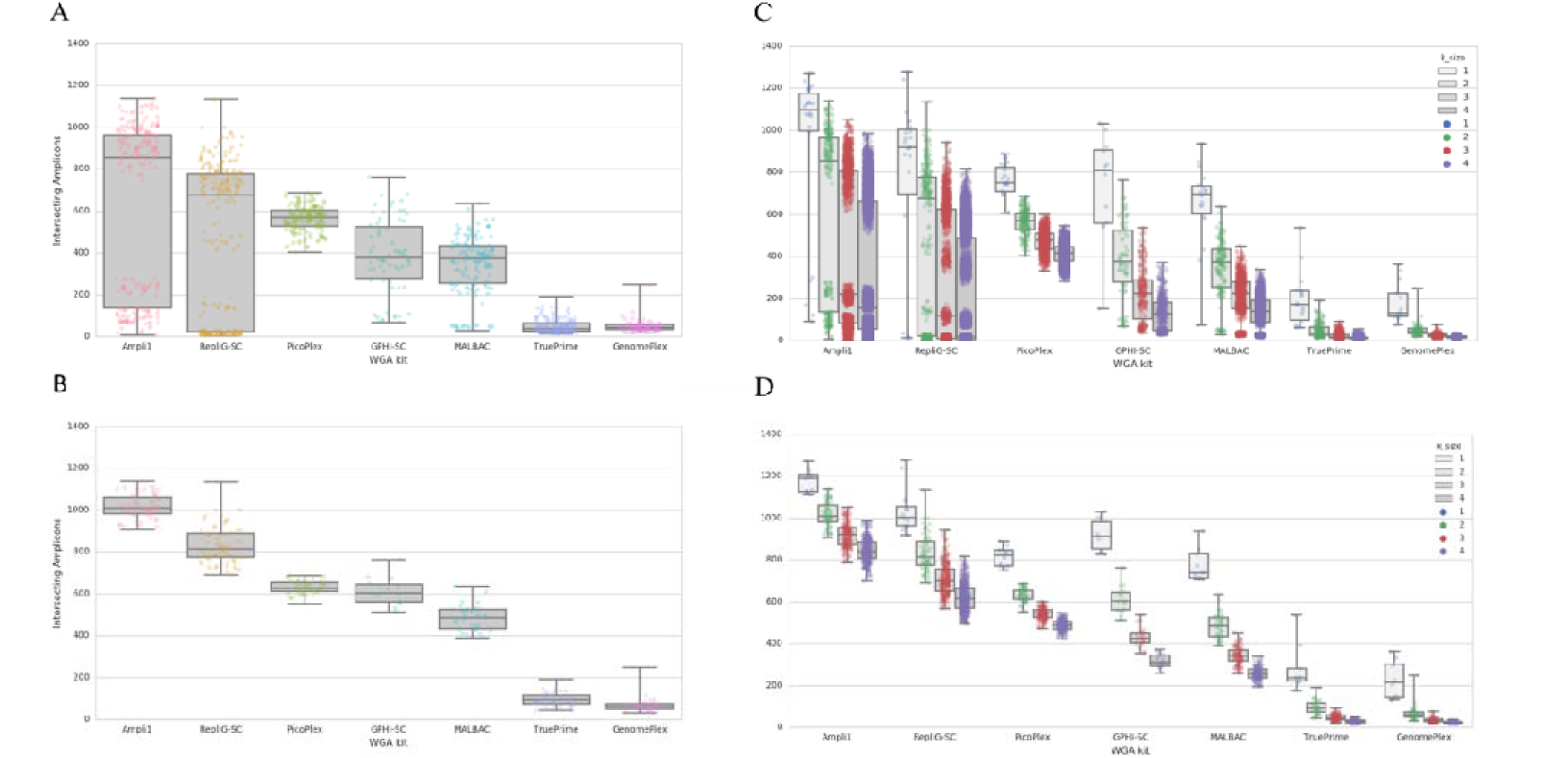
scWGA reproducibility analysis. A. Each pair of cells (k=2) per scWGA kit were analyzed for the number of mapped amplicons that were mapped in both cells. Each dot represents a pair of cells, y-axis is the number of amplicons which were mapped by >0 reads that worked for both cells (intersecting amplicons). B. Same analysis as in (A) but for the upper median of the most successful cells as reflected by their number of amplifying amplicons (Figure 1). C,D. Same analysis as in A and B, respectively for cell groups comprised of k=1 to k=4 cells (k=1 is the same as presented in Figure 1, k=2 is the same as presented inA and B. Repeated presentation is for visualization of results in the context of all group sizes).

### scWGA error rate analysis

scWGA template is a single cell genome, which essentially is single copy of every nucleotide (besides specific cell cycle periods that increase the number of copies to 2, resulting in improved ADO ratio for scWGA products^15^). Therefore, any *in vitro*mutation insertion, specifically at early stages of amplification, may lead to untraceable mutations that eventually are genotyped as real data. We opted to generate a numerical grading of different kits by using the intrinsic nature of MS loci which makes them mutation prone both *in vivo*and both *in vitro*^14^. We used the sequencing results and our lab’s MS genotyping tool to generates a simulated stutter noise score for each MS target^16^: In summary, sinceMS naturally undergo *in vitro* noise insertion during amplification, the repeat number histogram of their sequenced reads reflects this mutational processes. The MS genotyping tool follows these mutational processes by simulation, resulting in not only a genotyping answer but also a confidence score (Supplementary Figure 3) and an estimated amount of amplification the sample undergone to which the confidence is orthogonal. Since the downstream *in vitro*amplification is equal for every WGA sample, the noise difference between different kits is trailed from the scWGA reaction itself, and hence can be used as comparative measure to determine the differences between the error-rate per kit, as reflected by thousands of analyzed loci. For the error rate measurement, we have analyzed only AC type MS loci, with more than 30 reads, from X chromosome,to get a clear mono-allelic signal. We than plotted all estimated amounts of amplification for the all loci X cells in each WGA kit in a comparative plot (Figure 3). As a general control, the expected outcome should present an advantage of MDA based methods over other PCR based methods, as we see for RepliG-SC (GPHI-SC, which is also MDA based follows, together with GenomePlex).

**Figure 3.**
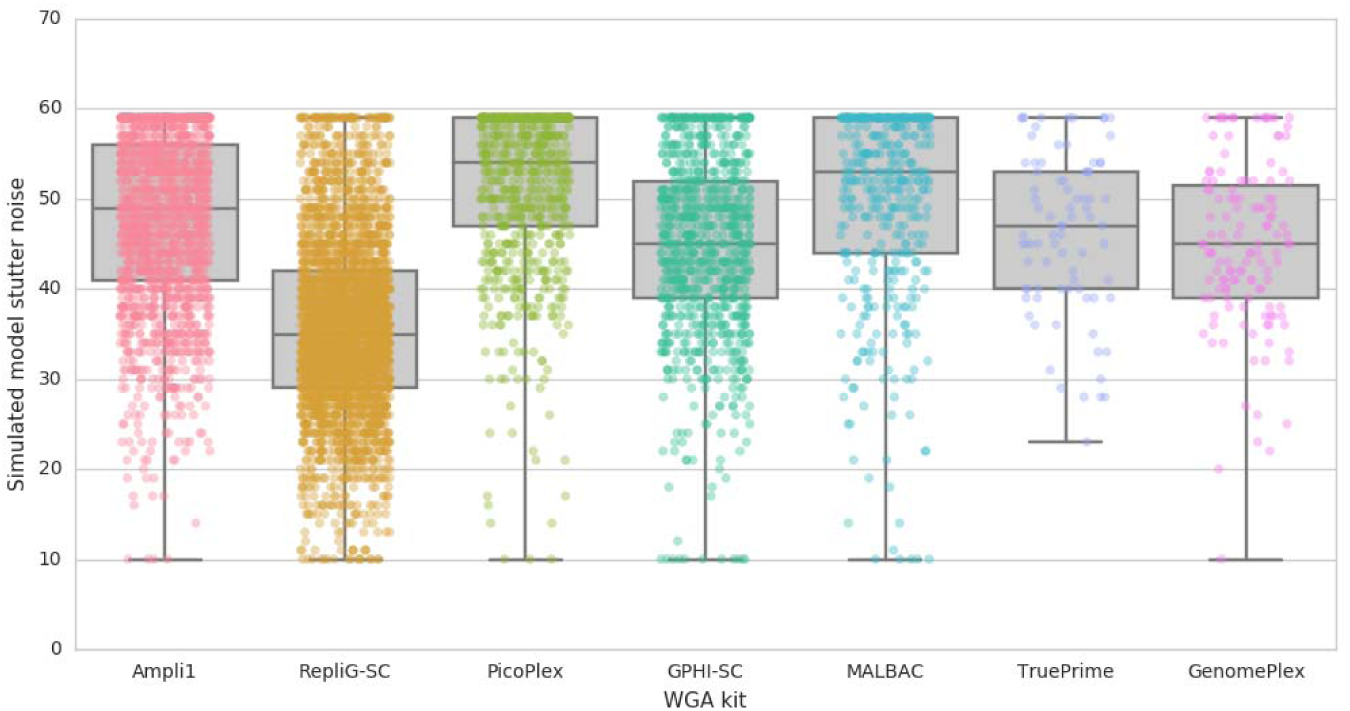
Error rate analysis of different scWGA kits. AC type MS loci targets from the X chromosome (>30 reads) were analyzed using our MS genotyping tool^16^ and estimated amount of amplification was fitted by simulation (Y axis, lower is better). All analyzed loci X cells per each kit were plotted to enable a comparison between different scWGA error rates.

## Discussion

In recent years, due to development of sequencing technologies and the rise of the single cell genomics field, there is an increased demand for scWGA kits that are accurate and robust^1^. The number of applications for SC genomic analysis is extreme, but the main interest in SC genome analysis is coming from the cancer research, as there is a growing interest in the variance between cells within the cancer cell population, which requires the accurate analysis of its individual composing cells^17^. Single cell experiment usually requires tens to thousands of cells, and hence comparing several cells per kit is not comparable to a real experiment. In this study, we opted to conduct the largest scale scWGA kit comparison, containing, for the first time all currently available commercial scWGA kits (that the authors knew at the time of the experimental design). Notably, improvements on existing kits were developed, namely upon MDA that is simpler than PCR based methods, as it has less processing steps and is based upon isothermal amplification. These modifications require specific equipment^18,19^ or experimental design^15^ (cell stage or limited amplification). Therfore, to enable a comparison dedicated to wide research community, we chose only commercially available kits and followed their exact manuals (or adjusted in accordance with the manufacture guidelines, see methods).

To this end we have utilized our established automated targeted enrichment based protocol^14^ on 125 different SCs. After normalizing the total reads per sample (and hence per kit), we sequenced and measured of different scWGA comparative categories: genome coverage, reliability, reproducibility and error rate.

Taking Ampli1’s internal bias against MseI containing amplicons (Supplementary Figure 2) into account, Ampli1 is the best kit in coverage and reproducibility. This biased amplification should be considered when planning an experiment. In most cases, one can order targeting probes/ primers that consider Ampli1 biased amplification; however, this kit may not be suitable for several application types. The reproducibility property of a kit should is sometime more important than of genome coverage.

Phylogenetics algorithms compare same data points (e.g. loci, SNPs) in every analyzed sample and later generate trees that reflect that comparison. When analyzing single cells for their cell lineage relationship^4^, ADO plays an important drawback in such cell lineage tree reconstruction experiments, as it reduces the number of analyzed loci^20^. However, in such phylogenetics algorithms the comparable number of loci effect is even larger that the successful coverage per cell. For example: a data set of 70% genome coverage for two cells can range between fully reproducible loci number (70% of the data is comparable) to low reproducibility (until 40% comparable loci). In summary, reproducibility makes a kit more suitable for single cell lineage analysis experiments^20^.

In this experiment, PicoPlex was proved to be the most reliable kit, showing repeated results forall cells, both in the coverage perspective and both in reproducibility perspective, with low variance for all analyzed cells (Figure 2a and Figure 2c). We chose to also present the data of the upper median of the most successful cells (Figure 2b and Figure 2d) as a simulation of a real experiment, where the best cells are chosen for analysis. In specific cases, such as with rare cells populations, this is selection is not an option. Moreover, the high cost per sample, ranging between 15$-36$ per cell, makes the reliability improvement a key cost factor for a large-scale experiment. We believe that a fine calibration of every step of an experiment, from the cell picking, to the WGA procedure can achieve improved results that improve the reliability factor. Results show that GenomePlex and TruePrime did not work as well as other kits in our hands. The random distribution of samples in the AA chips rules out the batch effect reason for their failure, in addition to the fact that most kits also underwent the cell picking and WGA procedure at the same day (see Methods). Positive controls of both GenomePlex and TruePrime provide more or less the same amplicon count results as their best single cells, suggesting that either the kit poorly worked in our hands or that its reaction output is biased in our targeted enrichment panel. We suggest that further calibrations of their protocol may improve their results. One can also suggest that the best two kits did not undergo purifications. Again, we followed the recommendations of the manufacturers and further calibration may improve the results; however, for both of the above arguments, kit calibration was not in the scope of this research, as it specifically validated the success per kit according to its recommended off-the-shelf protocol using our targeted enrichment assay.

The current methodology to track and compare scWGA error rate is by comparing the sequencing data generated from scWGA products to a reference genome, and therefore relies on a prior knowledge, which in the case of MS can be prone to errors or even not exist. We used a mapping approach that takes into account all the possible repeat numbers a MS can have to avoid mapping bias and employed our lab’s dedicated genotyping tools^16^. As expected RepliG-SC wins as it is based on isothermal amplification that was described as having a low error rate than other WGA protocols^3^. This makes it favorable for specific applications such as indel/single nucleotide variations, which are sensitive for the polymerase error rate, specifically in SC experiments. GPHI-SC and TruePrime, another MDA based methods are amongst the three following kits, together with GenomePlex. Nevertheless, both TruePrime and GenomePlex have much less data points, due to their low success in this experiment.

Although the starting template for the AA chips are WGA product, not normalized for their concentration, all of the kits manufacturers declare that the yield per single cell is micrograms to tens of micrograms (1 fold difference). Since every PCR process yields sufficient amplification that presumably reaches a plateau (also visible in the second PCR using real time amplification), the difference between the amplification cycles per kit should be of maximum 3-4 cycles. The presented data on Figure 3 simulated the number of noisy amplification cycles per kit. Even after adding the possible 3-4 amplification cycles to RepliG-SC (assuming its samples underwent less amplification cycles due to its higher concentration), it is still the best kit in the error rate aspect, as expected from isothermal amplification^3^.

Critical points that may be raised towards this comparison are (1) **The experiment is limited to a targeted enrichment panel and is not a true random WGS experiment.**We acknowledge that it is not a random experiment and the use of targeted enrichment as a subset of the genome is not equivalent to WGS. In addition, in previous comparisons it was shown that with a low coverage per sample one could detect the coverage at a high significance^9^. However, even at a low depth of coverage, the cost per genome is not scalable to simulate a real SC experiment that usually comprises of tens to thousands of analyzed cells. The use of targeted sequencing therefore offers a cheap and reliable measurement that mimics a real experiment. Our experimental design advantages are: (a) cheap and therefore enables a large examined cohort of cells per kit, (b) comprises of a large amplicon panel (3401 amplicons) for improved statistics, and (c) relies on a genome template of a diploid normal cell line. This cell, when analyzed for X chromosome only, yields clear results that are less noisy than other cell types. Importantly, our results are in agreement with the common literature (namely MDA effectiveness in genome coverage and error rate) and therefore present a powerful and cost-effective tool for scWGA kit comparison assay. (2) **The panel is mostly comprised of MS containing amplicons**that may bias the probability of amplification and affect the conclusions of the genome coverage and reproducibility results. A biased effect of this kind would result in a change inthe composition of read counts per sample (MS containing amplicons and non-MS containing amplicons),compared with the original panel composition. To rule our this option we examined the above amplicon count composition of H1 bulk templates (duplicates) and compared them to the composition of the original panel. While the original panel composition is 95% and 5% (MS containing amplicons and non-MS containing amplicons, respectively), the compositions of amplicons count of H1 bulk template duplicate are 95.3% and 4.7% for duplicate 1, and 95.4% and 4.6% for duplicate 2, respectively. This proves that the amplicon count was not biased by amplicon composition. (3) **One can choose to increase the panel size to improve statistics (*e.g.* exome panel).**Increasing the probe/primers panel to larger genome panel will probably enable better statistical analysis.

However, this will also dramatically affect the cost as in most cases, a change in targeted enrichment protocol will be required, and the cost per sequencing (of more bases) will also be increased.

We acknowledge that the targeted enrichment assay has its limitations and is not suitable for all analysis types. Analysis of amplification uniformity is not accurate using this assay as it is basedon PCR, which is template sensitive, making its read coverage less informative for accurate analysisof *e.g.* copy-number profiling. In addition, chimaeras, artefact joining of two separated genomic regions is overlooked in our assay. Affiliated to MDA based analysis, chimaeras will not be detected as it will either not be amplified (if amplicon was not joined as a whole) or will be amplified without knowing that it happened (if a chimera happened by joining of the entire amplicons).

It is clear from previous scWGA kit comparison experiments and from the presented comparison that there is not a single winner in the race for the best scWGA kits, but several exceling kits, depending on the category of interest. Overall, this comparative assay demonstrates a cost-effective benchmark to compare different WGA kit properties of analyzed SCs and enables an educated selection of a WGA of choice, depending on the type of required analysis.

## Methods

### Generation of clonal human ES cells

H1 human ES cells (WA01) were obtained from the WiCell Research Institute (Madison, WI). In order to create single cell (SC) clones, SCs were picked and deposited in separated single 96 wells using an automated cell picking device (CellCelector, ALS). Cells were cultured and treated as described in ^14^.

### Cell Picking and scWGA amplification

At the day of the experiment, before cell treatment, in order to minimize contamination, a clean working environment was established: bench and pipettes were RNase and DNA decontaminated (RNase AWAY, Molecular BioProducts) and consumables (PCR plates, tubes, tips etc.), non-kit reagents (PCR grade water, buffers etc.) and pipettes were UV irradiated.

Followed by ~2 weeks of growth, a single clone was selected and detached to enable selection and picking of single cells. For the cell picking procedure cells were treated as described in^14^with the exception that prior to picking to scWGA reactions, cells were detached and dissociated using 550 U/ml StemPro Accutase Cell Dissociation Reagent (Gibco), incubated for 6 minutes, pulled down, re-suspended in iMEFs Conditioned hESC medium (CM) and were spread over a 6cm non-adherent petri dish.

Individual cells were picked and deposited in separated single wells in dedicated 96-well PCR plates (Axigen, each kit in a separate plate), containing the required reagent. All pickings were performed on the same clone and at the same day (besides GPHI-SC, which was performed a few days later due to technical reasons). 1μl of positive (100pg Hela genomic DNA, NEB) and negative (water) controls were added as templates to non-deposited reaction wells. Samples were directly processed by their appropriate scWGA protocol. All reagents were immediately or sequentially (depending on the protocol) added to the original picking well, without material transfer, to avoid material loss.

Reagents, volumes, procedures and cleanups (if applicable) were as described in the scWGA protocols (see also list in Table1) with the following modifications, which were suggested and recommended by the manufactures to fit a >2μl deposition volume: (1) Ampli1 - The deposition volume was changed to 5μl PBS and although the kit was Version 2, Version 1 was performed (similar to Version 2 but with a 3-day protocol). (2) TruePrime - The deposition volume was changed to 5μl PBS, lysis was performed at 65°C, amplification mix contained 19.3μl water, followed by a 3-hour reaction. (3) GPHI-SC - The deposition volume was changed to 4μl water. 1μl lysis buffer (DTT 250mM, KOH 1M, Tween20 0.05%) was added to the reaction. Following lysis, 2.5μl of Neutralization buffer were added. Amplification mix (composed of: 1.5μl Enzyme Mix, 3μl water and 16.5μl Reaction Buffer) was added to each reaction, followed by 2 hours of amplification at 30°C and 65°C inactivation for 10 minutes.

An H1 cell bulk extraction (DNeasy Blood & Tissue Kit, Qiagen) was added as another positive control to the experiment.

### Processing DNA samples in the cell lineage analysis platform

We have utilized our lab developed cell lineage analysis platform as previously elaborated in^14^ to generate targeted enrichment NGS data from every cell sample. In summary, the cell lineage analysis platform is a pipelined, automated workflow that enables a large scale targeted enrichment analysis from many DNA samples. scWGA samples, their controls and the H1 bulk sample (in duplicate) were randomized and placed in an AccessArray (AA) chip (Fluidigm) for targeted enrichment using 3401 primer pairs, divided to 48 multiplex reactions (See Supplementary Table 1 for description of primers and multiplex groups and Supplementary Table 2 for description of all MS loci in the panel). Positive and negative controls (HeLa Genomic DNA 100ng/μl & water, respectively) were added as additional controls per each of the 5 AA. AA PCR protocol was as in^14^. Barcoding PCR was modified from the original protocol to the NEBNext Ultra II Q5 Master Mix (M0544, NEB) protocol, with 0.5μM primer concentration and addition of SYBR green I (Lonza) at a final reaction of 0.5X. Samples were purified and sequenced (Miseq, Illumina) after pooling in in an equal volume per sample (performed by Echo550 liquid handler robot, Labcyte). Total reads analysis was used to normalize each sample volume to yield as equal as possible read count per sample, and as a result, per kit, using the Echo550 robot, Deep-sequencing was later performed (NextSeq, Illumina). Samples that did not pass a reads threshold were also pooled: samples with >60% mapped reads were pooled at a fix volume of 6.5μl, samples with <60% and negative controls were pooled at the average volume of the normalized samples (667.5nl).

### Computational analysis

The MS-aware mapping of next generation sequencing was done using the pipeline described in Biezuner *el al*^14^. MS mutation calling from repeat-number histograms was performed using the MS genotyping tool by Raz *el al*^16^. The error rate analysis is based on AC type MS from X chromosome, with >30 reads.

## Data access

Sequencing data generated in this study have been submitted to ArrayExpress (www.ebi.ac.uk/arrayexpress) under accession number E-MTAB-5968.

## Acknowledgments

We thank Dr. D. Pilzer for performing Access Array runs; and G. Cohen and Dr. H.M. Barr for operating the Echo550 robot runs.

This research was supported by the following foundations: The European Union FP7-ERC-AdG (233047), The EU-H2020- ERC-AdG (670535), The DFG (611042), The Israeli Science Foundation (ISF, P14587), The ISF-BROAD (P15439), The NIH (VUMC 38347) and The Kenneth and Sally Leafman Appelbaum Discovery Fund. Ehud Shapiro is the Incumbent of The Harry Weinrebe Professorial Chair of Computer Science and Biology.

